# Explicit control of step timing during split-belt walking reveals interdependent recalibration of movements in space and time

**DOI:** 10.1101/614644

**Authors:** Marcela Gonzalez-Rubio, Nicolas F. Velasquez, Gelsy Torres-Oviedo

## Abstract

Split-belt treadmills that move the legs at different speeds are thought to update internal representations of the environment, such that this novel condition generates a new locomotor pattern with distinct spatio-temporal features compared to those of regular walking. It is unclear the degree to which such recalibration of movements in the spatial and temporal domains is interdependent. In this study, we explicitly altered subjects’ limb motion in either space or time during split-belt walking to determine its impact on the adaptation of the other domain. Interestingly, we observed that motor adaptation in the spatial domain was susceptible to altering the temporal domain, whereas motor adaptation in the temporal domain was resilient to modifying the spatial domain. This nonreciprocal relation suggests a hierarchical organization such that the control of timing in locomotion has an effect on the control of limb position. This is of translational interest because clinical populations often have a greater deficit in one domain compared to the other. Our results suggest that explicit changes to temporal deficits cannot occur without modifying the spatial control of the limb.

## 1 INTRODUCTION

We are constantly adapting our movements to demands imposed by changes in the environment or our body. In walking, this requires the adaptation of spatial and temporal gait features to control “where” and “when” we step, respectively. Particularly, in split-belt walking when one leg moves faster than the other, it has been observed that subjects minimize spatial and temporal asymmetries by adopting motor patterns specific to the split environment (e.g., Malone et al., 2012). It is thought that this is achieved by updating internal representations of the treadmill for the control of the limb in space and time (Malone et al., 2012). There is a clinical interest in understanding the interdependence in the control of these two aspects of movement because pathological gait often has a greater deficiency in one domain compared to the other (Finley et al., 2015; Malone and Bastian, 2014). Thus, there is a translational interest to determine if spatial and temporal asymmetries in clinical populations can be targeted and treated independently.

Ample evidence supports that the adaptation, and hence control, of spatial and temporal gait features is dissociable. Notably, studies have shown that inter-limb measures such as step timing (temporal) and step position (spatial) adapt at different rates (Sombric et al., 2017; Malone and Bastian, 2010), they exhibit different generalization patterns (Torres-Oviedo and Bastian, 2010), and follow distinct adaptation dynamics throughout development (Vasudevan et al., 2011; Patrick et al., 2014) or healthy aging (Sombric et al., 2017). In addition, several behavioral studies show that subjects’ adjustment of spatial metrics can be altered (Malone and Bastian, 2010; Malone et al., 2012; Long et al., 2016) without modifying the adaptation of temporal gait features. However, the opposite has not been demonstrated. For example, altering intra-limb measures (i.e., characterizing single leg motion) of timing such as stance time duration (Afzal et al., 2015; Krishnan et al., 2016) also leads to changes in intra-limb spatial features such as stride lengths. In sum, the spatial and temporal control of the limb is thought to be dissociable, but it remains unclear if the adaptation of internal representations of timing can be altered and what is the impact of such manipulation in the temporal domain on the spatial control of the limb.

In this study we aimed to determine the interdependence between the spatial and temporal control of the limbs during walking, particularly of inter-limb parameters characterizing bipedal coordination. We hypothesized that spatial and temporal inter-limb features are adapted independently based on previous studies demonstrating their dissociation. To test this hypothesis, subjects walked on a split-belt treadmill, which requires the adaptation of spatial and temporal inter-limb coordination. We further altered subjects’ movements during split-belt walking by either instructing them “where” (spatial feedback) or “when” (temporal feedback) to take a step. We contrasted the impact of explicitly manipulating movements in one domain on the adaptation of the other domain to determine their interdependence.

## 2 MATERIAL AND METHODS

We recruited twenty-one healthy young subjects (13 women, 8 men, mean age 24.69 ± 4 years) to voluntarily participate in this study. Subjects were randomly assigned to three groups (n=7, each): 1) control, 2) spatial feedback, 3) temporal feedback to determine if explicitly altering the limb motion on either the spatial or the temporal domain with visual feedback during split-belt walking had an impact on the adaptation of the other domain (Figure 1A). Notably, if the control of these two domains was dissociable, altering one would not have an effect on the other. Alternatively, if they were interdependent, modifying the adaptation of one domain not only would have an effect on the targeted domain, but will also alter the other one. The protocol was approved by the Institutional Review Board of the University of Pittsburgh and all subjects gave informed consent prior to testing.

**Figure 1:**
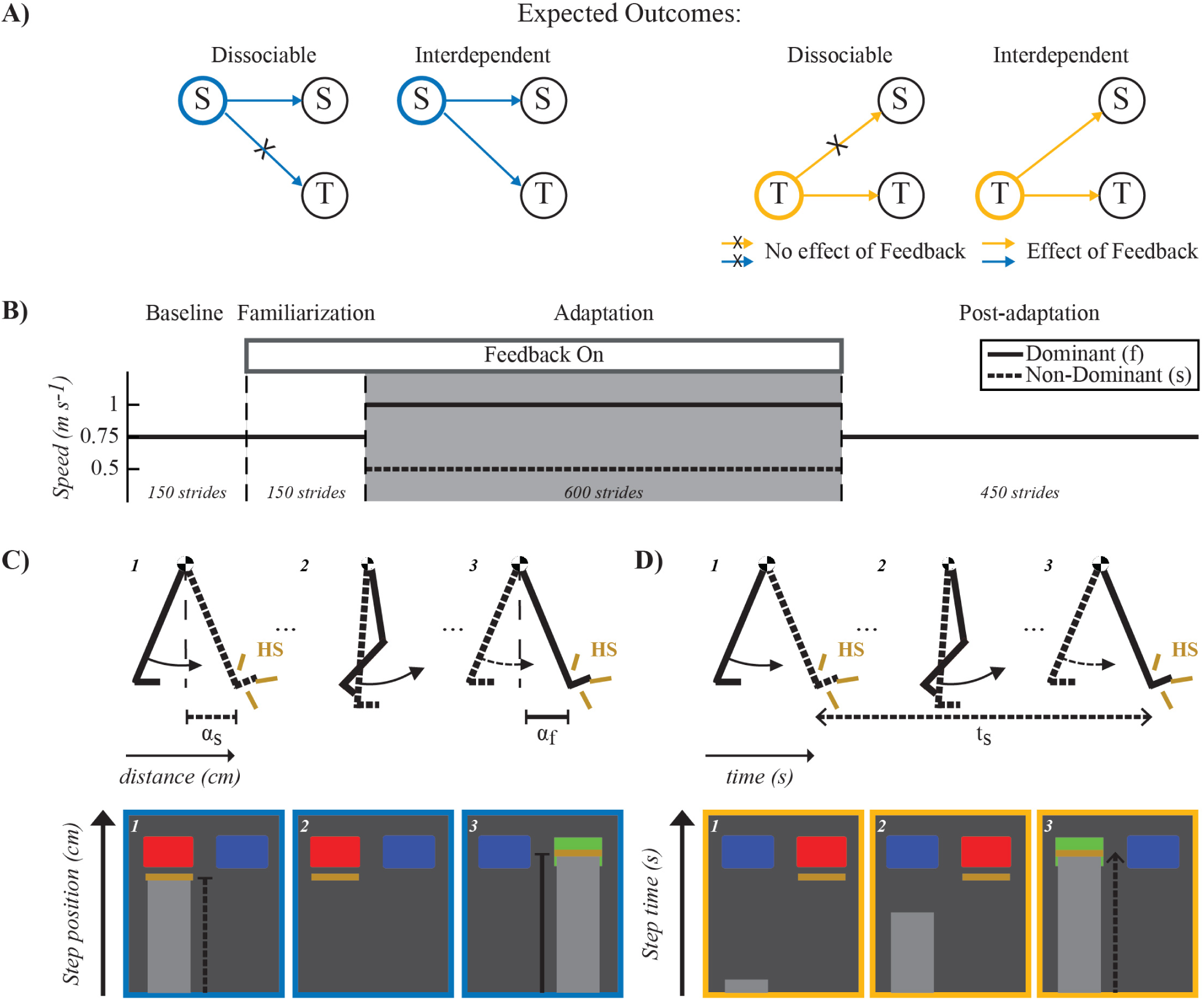
Expected outcomes, Paradigm and Feedback Visualization. **(A)** Expected outcomes for dissociable and interdependent internal representations of space and time. If dissociable, the feedback manipulation will only affect the targeted domain without changing the other domain. For example, spatial feedback (indicated with blue outline) would alter spatial features (S) of the motor pattern while temporal ones (T) remain invariant. On the other hand, if the domains are interdependent, feedback manipulation of one domain will also alter the other domain. For example, spatial feedback modifying spatial features of the motor pattern would also change temporal ones. **(B)** Split-belt walking paradigm used in all groups. Dashed lines separate the different experimental phases. All groups experienced the same number of strides during each phase (Baseline: 150, Familiarization: 150, Adaptation: 600, and Post-adaptation: 450). The two belts moved at the same speed (0.75*m/s*) during the Baseline and Familiarization phases. Only subjects in the feedback groups walked while observing their movements on a TV screen placed directly in front of them (Feedback On) during the familiarization phase. The feedback to these groups was also given during the Adaptation phase (gray shaded area) during which one belt (fast belt) moved at 1*m/s* and the other one (slow belt) moved at 0.5*m/s*. Finally, during Post-adaptation subjects walked again with the two belts moving at the same speed (0.75*m/s*). **(C-D)** Visual feedback schematic. Schematic of the legs in the top row illustrate the step position (e.g., *α*_*f*_ and *α*_*s*_) and step time (e.g., *t*_*s*_), which were the walking features used in the spatial and temporal feedback tasks, respectively. Bottom rows in panel C and D illustrate the screen shots observed by individuals in the spatial feedback group (Panel C) or in the temporal feedback group (Panel D). Blue rectangles indicated the target step position or step time value that subjects had to achieve with each leg. These rectangles turned green when subjects met the desired step position or step time values and red when they did not. Yellow lines indicated either the step position value (Panel C) or the step time value (Panel D) at heel strike (HS) when taking a step with the right or left leg (e.g., left leg’s step position is shown in the screen shot #1). In the example shown, the step position was correct for the right leg but not for the left leg. The light grey progression bars showed in real-time either the the distance from the ankle to the hip markers as subjects swing the leg forward (Panel C) or the time that the subject had spent on the standing leg since it hit the ground (Panel D).

### 2.1 Experimental Protocol

All subjects walked on a split-belt treadmill during four experimental phases: Baseline, Familiarization, Adaptation, and Post-adaptation. The speed for each belt during these phases is shown in Figure 1B. This speed profile enabled individuals to walk at an averaged speed of 0.75 m/s throughout the experiment. In the Baseline phase, individuals walked with the two belts moving at the same speed of 0.75 m/s for 150 strides (∼3 min). Recordings from these phase were used as the reference gait for every individual. In the Familiarization phase, all participants also walked at 0.75 m/s for 150 strides, but only subjects in the feedback groups received the same visual feedback that they were going to experience during the subsequent Adaptation phase. This was done to allow feedback groups to become habituated to use the provided visual feedback to control either spatial (spatial feedback group) or temporal (temporal feedback group) gait features. In the Adaptation phase, the belts were moved at a 2:1 ratio (1:0.5 m/s) for 600 strides (∼13 min). We selected these specific belt speeds because other studies have indicated that they induce robust sensorimotor adaptation (Reisman et al., 2005; Mawase et al., 2014; Sombric et al., 2017; Vervoort et al., 2019) and we observed in pilot tests that subjects with visual feedback at these speeds could successfully modify the spatial and temporal gait features of interest. The self-reported dominant leg walked on the fast belt. In the Post-adaptation phase, all individuals walked with both belts moving at 0.75 m/s for 450 strides (∼10 min). This phase was used to quantify gait changes following the Adaptation phase. The treadmill belts were stopped at the end of each experimental phase. A handrail was placed in front of the treadmill for safety purposes, but individuals did not hold it while walking. A custom-built divider was placed in the middle of the treadmill during the entire experimental protocol to prevent subjects from stepping on the same belt with both legs. Subjects also wore a safety harness (SoloStep, SD) that did not interfere with their walking (no body weight support).

We tested three groups: 1) control group, 2) spatial feedback group, 3) temporal feedback group. The control group was asked to “just walk” without any specific feedback on subjects’ movements. Each subject in the spatial or temporal feedback groups was instructed to either maintain his/her averaged baseline step position (spatial feedback group) or averaged baseline step time (temporal feedback group) when the feedback was on. Step position was defined as the sagittal distance between the leading leg’s ankle to the hip at heel strike (Figure 1C). Step time was defined as the time period from heel strike (i.e., foot landing) of one leg to heel strike of the other leg (Figure 1D). We chose to manipulate step position and step time for consistency with other studies (Malone et al., 2012; Long et al., 2016) and because these parameters are adjusted during split-belt walking to reduce spatial and temporal inter-limb asymmetries, respectively (Malone et al., 2012). Panels C and D in Figure 1 show sample screen shots of the visual feedback observed by each group on a screen placed in front of them. More specifically, we permanently displayed either spatial or temporal targets (blue rectangles) indicating the averaged step position (spatial feedback group) or averaged step time (temporal feedback group) across legs during baseline walking. These targets turned green when subjects achieved the targeted baseline values and they turned red when they did not. A tolerance of ±0.75% and ±1.25% of the baseline value was given to subjects in the spatial and temporal feedback groups, respectively. Yellow lines indicated the actual step position and step time for each leg at every step. Thus, subjects could appreciate how far they were from the targeted spatial or temporal value at every step.

### 2.2 Data Collection

Kinetic and kinematic data were collected to quantify subjects’ gait. Kinematic data was collected at 100 Hz with a motion capture system (VICON motion systems, Oxford, UK). Passive reflective markers were placed bilaterally on bony landmarks at the ankle (malleolus) and the hip (greater trochanter). Kinetic data was collected at 1000 Hz with the instrumented split-belt treadmill (Bertec, OH). The normal ground reaction force (*F*_*z*_) was used to detect when the foot landed (i.e., heel strike) or was lifted off (i.e., toe off). A threshold of 10 N was used for detecting heel strikes and toe offs for data analysis, whereas a threshold of 30 N was used for counting strides in real-time.

### 2.3 Data Analysis

#### 2.3.1 Gait parameters

We computed six gait parameters previously used (Malone et al., 2012) to quantify the adaptation of spatial and temporal control of the limb during split-belt walking: *S*_*out*_, *T*_*out*_, *S*_*A*_, *T*_*A*_, *S*_*nA*_, and *T*_*nA*_. We used *S*_*out*_ and *T*_*out*_ because our feedback was designed to directly alter these metrics. For example, subjects in the spatial feedback group were given feedback to maintain the same baseline step position in both legs. *S*_*out*_ is, therefore, a good metric of performance for the spatial feedback group since it quantifies the difference in step positions, *α*_*f*_ and *α*_*s*_, when taking a step with the fast and slow leg, respectively. Formally expressed:

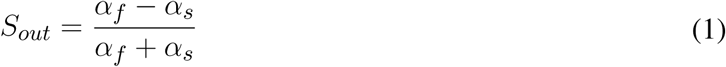

*α*_*i*_ is a length measurement that indicates the position of the ankle marker relative to the hip marker at heel strike. The subscript *i* can be either *f* or *s* for the leg that is on the fast belt or slow belt, respectively. By convention, *S*_*out*_ is positive when the fast leg’s foot lands farther away from the body when taking a step than the slow leg’s one (i.e., *α*_*f*_ *> α*_*s*_). *S*_*out*_ is zero during baseline and subjects in the feedback group were instructed to maintain this value during split-belt walking.

Similarly, subjects in the temporal feedback group were given feedback to maintain the same baseline step times in both legs. *T*_*out*_ is, therefore, a good metric of performance for the temporal feedback group since it quantifies the difference in step times, *t*_*s*_ and *t*_*f*_. Step time (*t*_*s*_) is defined as the time interval to take a step on the slow belt (i.e., duration from heel strike on the fast belt to the subsequent heel strike on the slow belt) and vice versa for *t*_*s*_. Formally expressed:

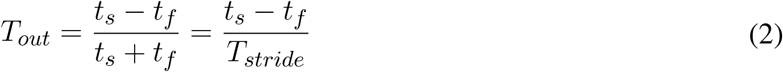

Where *T*_*stride*_ is the stride time (i.e., time interval between two consecutive heel strikes with the same leg). By convention, *T*_*out*_ is positive when the slow leg’s step time is longer that the fast leg’s one. *T*_*out*_ is zero during baseline and subjects in the feedback group were instructed to maintain this value during split-belt walking. It has been previously shown that *S*_*out*_ and *T*_*out*_ are adapted during split-belt walking to minimize spatial and temporal baseline asymmetries defined as *S*_*A*_ and *T*_*A*_, respectively (Malone et al., 2012). Therefore, we also quantified *S*_*A*_ and *T*_*A*_ because these are adaptive parameters (Malone et al., 2012; Reisman et al., 2005; Malone and Bastian, 2010) that could be indirectly altered by our spatial and temporal feedback even if subjects in these groups were not explicitly instructed to modify them.

*S*_*A*_ quantifies differences between the legs in where they oscillate with respect to the body. The oscillation of each leg was computed as the ratio between two distances: step position (*α*) and stride length (*γ*) (i.e., anterior-posterior distance from foot position at heel strike to ipsilateral foot position at toe off). Thus, *S*_*A*_ (legs’ orientation asymmetry) was computed as the difference between these ratios when taking a step with the slow leg (i.e., slow leg leading) vs. the fast leg (see Eq. 3).

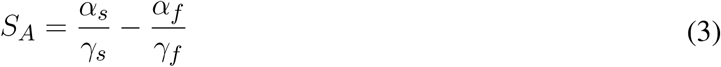

In the temporal domain, *T*_*A*_ quantified the difference in double support times (i.e., period during which both legs are on the ground) when taking a step with the fast leg (*DS*_*s*_) or slow leg (*DS*_*f*_), respectively (see Eq. 4). In other words, *DS*_*s*_ is defined as the time from fast heel strike to slow toe off and *DS*_*f*_ as the time from slow heel strike to fast toe off.

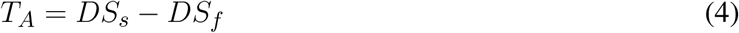

Lastly, we computed gait parameters defined as *S*_*nA*_ and *T*_*nA*_, to test the specificity of our feedback. Namely, it has been previously observed that these parameters do not change as subjects walk in the split-belt environment (Malone et al., 2012; Reisman et al., 2005; Yokoyama et al., 2018). Thus, these measures are thought to simply reflect the speed difference between the legs, and hence, we expected that our feedback would not alter them. Specifically, *S*_*nA*_ quantifies the difference between the fast and slow leg’s ranges of motion *γ*_*f*_ and *γ*_*s*_. Formally expressed as:

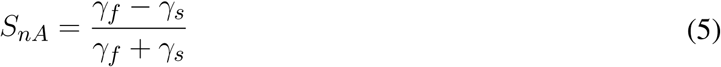

The non-adaptive measure in the temporal domain *T*_*nA*_ quantifies the difference between the slow and fast leg’s stance time durations (which is defined as the interval when the foot is in contact with the ground), which we labeled as *ST*_*s*_ and *ST*_*f*_, respectively. Formally expressed as:

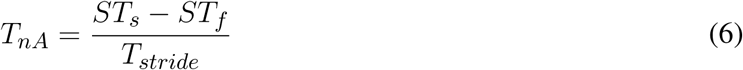

#### 2.3.2 Outcome measures

We computed *steady state* and *after-effects* to respectively characterize the adaptation and recalibration of walking in the spatial and temporal domains. Both of these outcome measures were computed for each gait parameter described in the previous section. *Steady state* was used to characterize the spatial and temporal features of the adapted motor pattern once subjects reached a plateau during split-belt walking. *Steady state* was computed as the averaged of the last 40 strides during the Adaptation phase, except for the very last 5 strides to exclude transient steps when subjects were told to hold on to the handrail prior to stopping the treadmill. *After-effects* were used to characterize the recalibration of subjects’ internal representation of the environment (Roemmich and Bastian, 2015) leading to gait changes that were sustained following split-belt walking compared to baseline spatial and temporal gait features. *After-effects* were computed as the averaged value for each gait parameter over the first thirty strides of post-adaptation. We used 30 strides, rather than only the initial 1 to 5 strides, because we were interested in characterizing long lasting after-effects (Long et al., 2015; Mawase et al., 2017; Roemmich and Bastian, 2015). We removed baseline biases from both measures by subtracting the baseline values for each gait parameter averaged over the last 40 strides during baseline (minus the very last transient 5 strides). This was done to exclude individual biases before aggregating subjects’ outcome measures in every group.

### 2.4 Statistical analysis

We performed separate two-way repeated measures ANOVAs (factors: group and epoch) comparing the control group to either the temporal or spatial feedback groups. This was done to determine the effect of experimentally altering either spatial or temporal measures during split-belt walking on outcome measures in both domains. When main effects of group or epoch were found (*p* < 0.05), we used Fisher’s LSD *post-hoc* testing to assess if main effects were driven by differences between the control group and feedback group in either domain. We applied a Bonferroni correction to account for 2 comparisons of interest resulting in a significance level set to *α* = 0.025. We selected to do our analysis with unbiased data (i.e., subject-specific baseline bias removed) to reduce inter-subject variability due to distinct baseline biases and focus on group effects due to the distinct experimental manipulations. Lastly, we performed independent sample t-tests to determine if *steady state* or *after-effects* were significantly different from baseline. We applied Bonferroni corrections to account for 4 comparisons of interest (baseline vs. steady state and baseline vs. after-effects for each of the experimentally targeted *S*_*out*_ and *T*_*out*_ parameters) setting the significance level to *α* = 0.0125. For all other parameters, we set the significance level to *α* = 0.025 to account for only 2 comparisons of interest (baseline vs. after-effects in the spatial and temporal domains). This was done since we were primarily interested in the impact of the experimental manipulation on the *after-effects* of the parameters that were not explicitly targeted with the visual feedback.

## 3 RESULTS

### Confirmation of results supporting dissociable representation of spatial and temporal walking features

Spatial and temporal gait features adapted and recalibrated independently when feedback was used to alter the spatial control of the limb. This is indicated by the group differences qualitatively observed in the *S*_*out*_’s time courses during Adaptation and Post-adaptation (left panel in Figure 2A and 2B, respectively) contrasting the overlapping time courses of *T*_*out*_ in the control group (red trace) and spatial feedback group (blue trace) (right panel in Figures 2A and 2B). Accordingly, we found a significant group effect on *S*_*out*_ (*p* = 0.0039), but not a group (*p* = 0.3748) or group by epoch interaction effect on *T*_*out*_ (*p* = 0.2293). *Post-hoc* analysis indicated that the spatial feedback reduced the steady state of *S*_*out*_ relative to the control group (*S* →*S* : *p* = 0.0021); such that the steady state values reached by the spatial feedback group were not significantly different from zero (*p* = 0.0481), whereas those of the control group differed from zero (*p* = 0.0004). This indicated that individuals in the spatial feedback group were able to maintain their baseline *S*_*out*_ values with the visual feedback on this metric. In contrast, the steady state values of *T*_*out*_ were significantly different from zero in both groups (control group: *p <* 0.0001; spatial feedback group: *p* = 0.0004). The dissociation between spatial and temporal control was also shown by the after-effects of *S*_*out*_ and *T*_*out*_ in the control vs. spatial feedback groups (Figure 2B). *Post-hoc* analysis indicated that the spatial feedback group had reduced after-effects of *S*_*out*_ compared to the control group (*S* → *S* : *p* = 0.0159) and that only the control group had after-effects different from zero (control group: *p* = 0.0003; spatial feedback group: *p* = 0.0164). Conversely, *T*_*out*_ was once again not qualitatively different between the groups and the after-effects were non-significantly different from zero on either group (control group: *p* = 0.4235; spatial feedback group: *p* = 0.1023). In sum, spatial feedback had a domain-specific effect: it altered the adaptation and recalibration of *S*_*out*_ (targeted spatial parameter) without modifying the adaptation and aftereffects of step time (*T*_*out*_).

**Figure 2:**
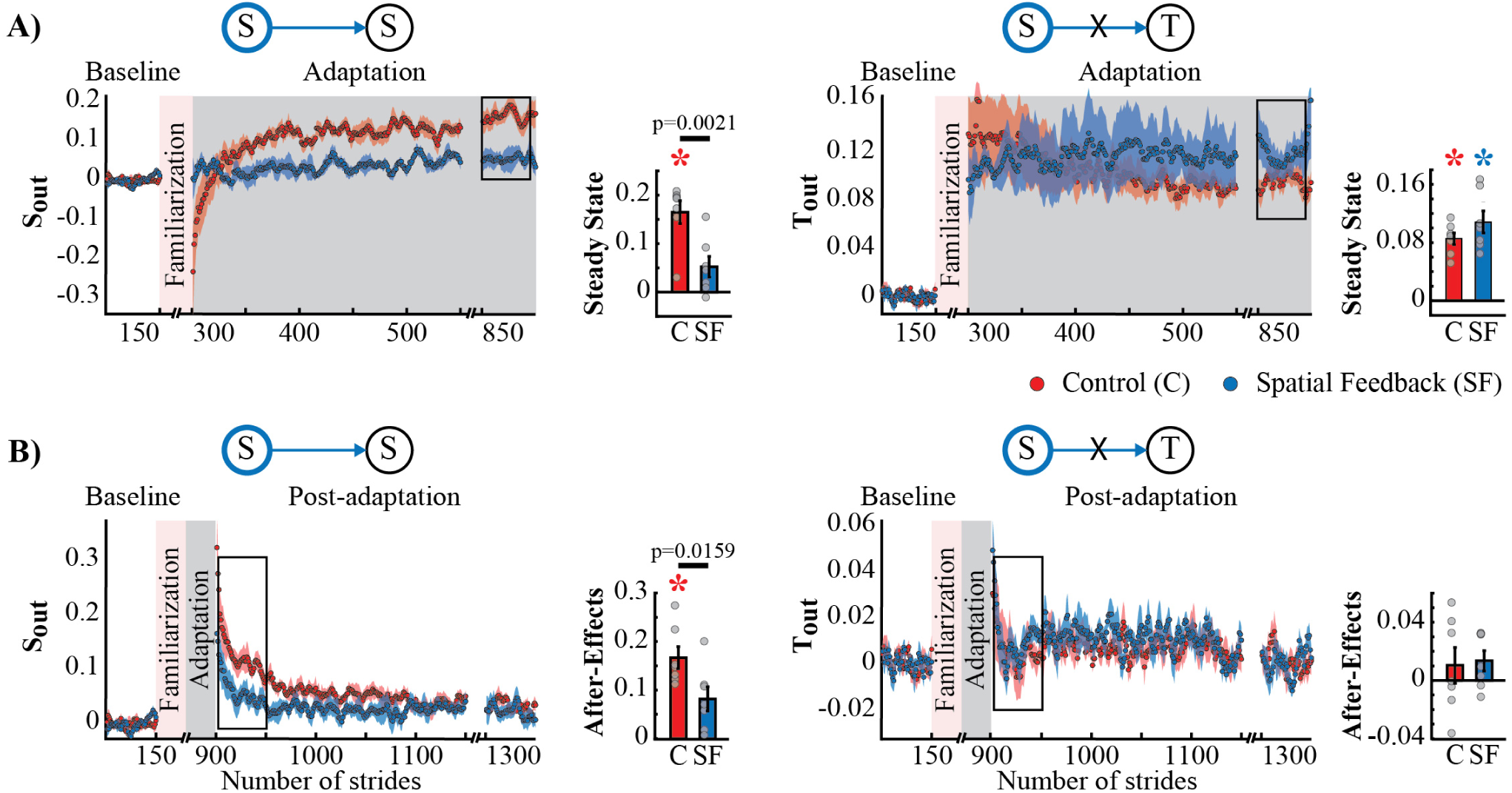
Adaptation and Post-adaptation of the parameters *S*_*out*_ (targeted) and *T*_*out*_ in the spatial feedback and control groups. Stride-by-stride time courses show the effect of altering step positions in the Adaptation (Panel A) and Post-adaptation (Panel B) of *S*_*out*_ and *T*_*out*_. Each data point in the time courses represents the average of five consecutive strides and shaded areas around the data points represent the standard errors. Bar plots indicate the mean average behavior in the epochs of interest (indicated with the black rectangles), gray dots indicate values for individual subjects, and vertical black lines are standard errors. Horizontal lines between bars illustrate significant differences between groups (*p* < 0.025). **A)** Steady state values of *S*_*out*_ and *T*_*out*_: We found a significant group difference in *S*_*out*_’s steady state. Colored asterisks indicate that the mean steady state for that group is significantly different from zero (*p* < 0.0125). **B)** After-effect values of *S*_*out*_ and *T*_*out*_: We found a significant group difference in *S*_*out*_’s after-effects. Colored asterisks indicate that the mean after-effect for that group is significantly different from zero (*p* < 0.0125).

The dissociation in adaptation and recalibration of spatial and temporal representations of walking was also supported by the analysis of spatial and temporal features known to be adapted by the split-belt task, but not directly targeted by our feedback. Namely, the spatial feedback also modified the Adaptation and Post-adaptation time courses of the legs’ orientation asymmetry quantified by *S*_*A*_, which is expected given its relation to *S*_*out*_. Note that the time courses of *S*_*A*_ for the spatial feedback group (blue trace) and control group (red trace) do not overlap during Adaptation and Post-adaptation (left panel Figure 3A and 3B). In contrast, the time courses of double support asymmetry (*T*_*A*_) were not altered by the spatial feedback, as shown by the overlap of *T*_*A*_ values during Adaptation and Post-adaptation of the temporal feedback and control groups (right panel Figure 3A and 3B). Consistently, we found a significant group effect in *S*_*A*_ (*p* = 0.0091) and a non-significant group (*p* = 0.8679) or group by epoch interaction (*p* = 0.2229) in *T*_*A*_. *Post-hoc* analyses revealed that between group differences in *S*_*A*_ were driven by the significantly different *S*_*A*_’s steady state (*S* → *S*_*A*_ : *p* = 0.0177) and trending differences in *S*_*A*_’s after-effects (*S* → *S*_*A*_ : *p* = 0.0358); such that after-effects were significant in the control group (*p* = 0.0009) but not in the spatial feedback group (*p* = 0.0542). Conversely, after-effects in double support asymmetry (*T*_*A*_) were significantly different from zero in all groups (control group:*p* = 0.0044; spatial feedback group:*p* = 0.0007). These results reiterated that changes in the spatial domain did not modify the temporal control of the limb in the temporal domain, replicating previous findings (Malone et al., 2012; Long et al., 2016).

**Figure 3:**
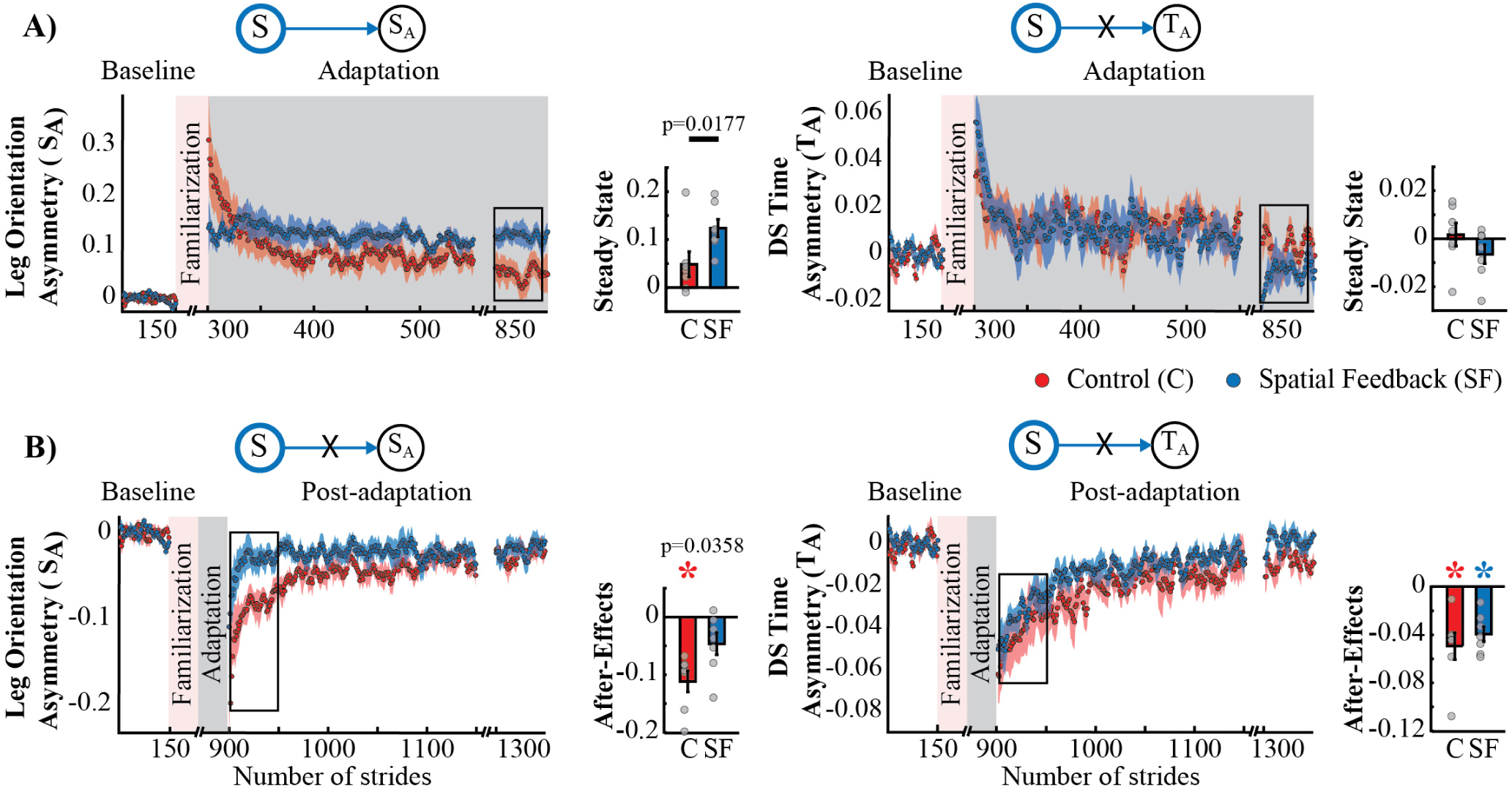
Adaptation and Post-adaptation for the adaptive but non-targeted parameters *S*_*A*_ (leg orientation asymmetry) and *T*_*A*_ (double support time asymmetry) in the spatial feedback and control groups. Stride-by-stride time courses show the effect of altering the step positions in the Adaptation (Panel A) and Pots-adaptation (Panel B) of *S*_*A*_ and *T*_*A*_. Each data point in the time courses represents the average of five consecutive strides and shaded areas around the data points represent the standard errors. Bar plots indicate the mean average behavior in the epochs of interest (indicated with the black rectangles), the gray dots indicate values for individual subjects, and vertical black lines are standard errors. Horizontal lines between bars illustrate significant differences between groups (*p* < 0.025). We found a significant group effect in *S*_*A*_. **A)** Steady States for *S*_*A*_ and *T*_*A*_: The significant group effect on *S*_*A*_ was driven by differences between the spatial feedback and control group in the non-targeted spatial motor output (adaptive motor output). **B)** After-Effects values of *S*_*A*_ and *T*_*A*_: We found significant group differences in *S*_*A*_. Colored asterisks indicate after-effect values are significantly different from zero (*p* < 0.025) according to *post-hoc* analysis.

### New evidence for interdependent representations of spatial and temporal walking features

Interestingly, we found that spatial and temporal gait features were not independent in their adaptation and recalibration when feedback was used to alter the temporal control of the limb. This is indicated by the qualitative differences between the time courses of *T*_*out*_ and *S*_*out*_ during the Adaptation (Figure 4A) and Post-adaptation phases (Figure 4B). Namely, the control group (red traces) and temporal feedback group (yellow traces) are different in both spatial and temporal parameters. Consistently, we found a significant group effect on *S*_*out*_ (*p* = 0.0005) and *T*_*out*_ (*p* = 0.0034). *Post-hoc* analyses revealed that the *T*_*out*_’s steady state was significantly different from zero in the control (*p* = 0.0004) and temporal feedback group (*p* = 0.0092). Thus, subjects in the temporal feedback group did not fully maintained the baseline values of *T*_*out*_, even if they were able to use the visual feedback to significantly reduce the *T*_*out*_ steady state during split-belt walking relative to the control group (*T* → *T* : *p <* 0.0001). While the temporal feedback group was designed to alter *T*_*out*_, we did not anticipate a reduction in the *S*_*out*_’s steady state relative to the control group (*T* → *S* : *p* = 0.0027) because this parameter was not directly targeted by the feedback. The interdependence between spatial and temporal domains was also shown by the analysis of after-effects in Post-adaptation (Figure 4B). *Post-hoc* analyses indicated that temporal feedback did not change the recalibration of *T*_*out*_ (*T* → *T* : *p* = 0.4663), but altered the recalibration of *S*_*out*_ (*T* → *S* : *p* = 0.0010). The non-significant effect on the recalibration of *T*_*out*_ was expected given that after-effects in this parameter are very short lived resulting in *T*_*out*_ after-effect values that are non-significantly different from zero (control group: *p* = 0.4235; temporal feedback group: *p* = 0.8550). In contrast, both groups had after-effects in *S*_*out*_ that were significantly different from zero (control group: *p* = 0.0003; temporal feedback group: *p* = 0.0021), but they were unexpectedly smaller in the temporal feedback group compared to the control group. In sum, the temporal feedback impact on adaptation and recalibration of *S*_*out*_ (spatial parameter) indicated an interdependence between the spatial and temporal control of the limb.

**Figure 4:**
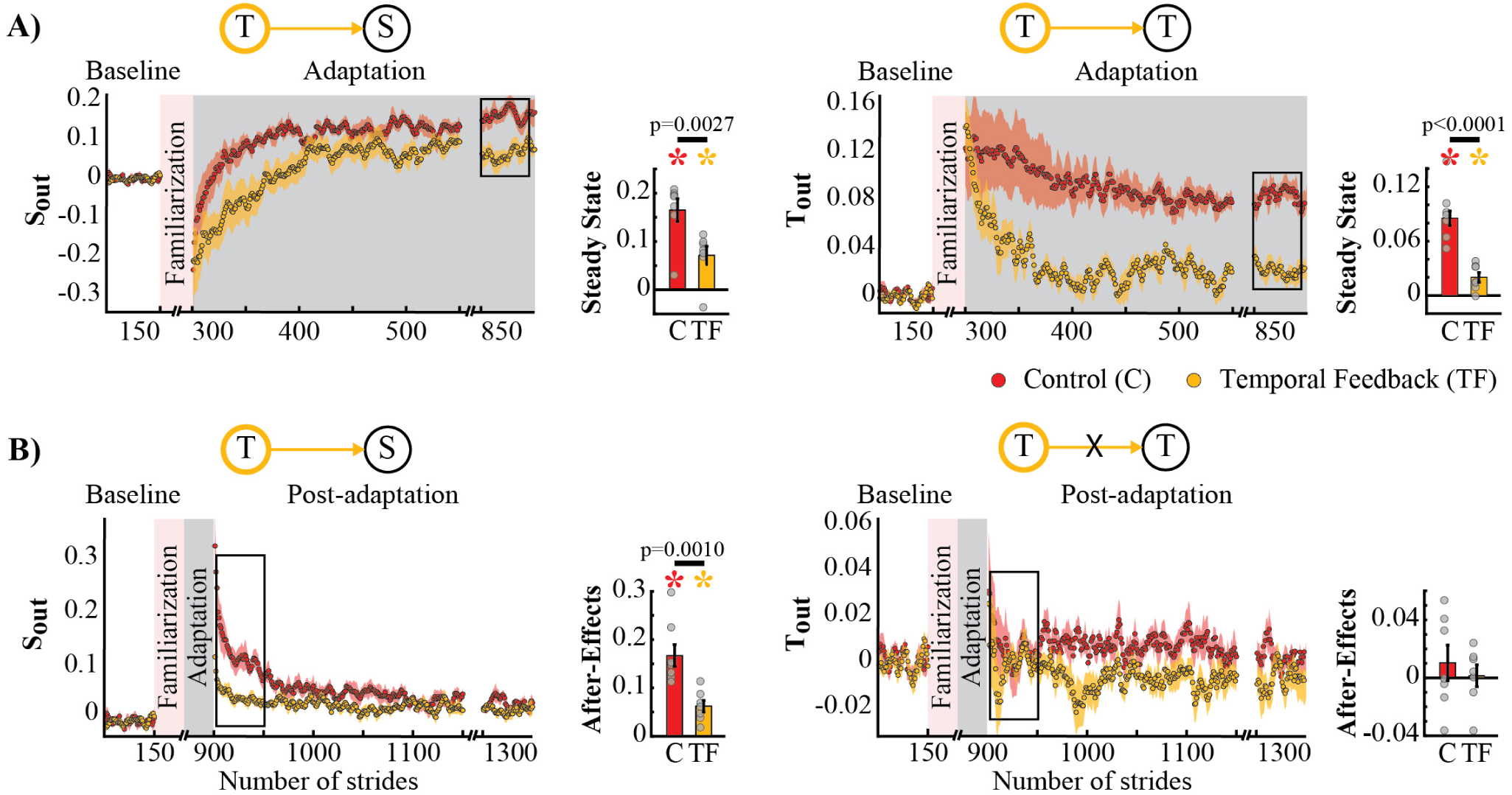
Adaptation and Post-adaptation of the parameters *S*_*out*_ and *T*_*out*_ (targeted) in the temporal feedback and control groups. Stride-by-stride time courses show the effect of altering step times in the Adaptation (Panel A) and Post-adaptation (Panel B) of *S*_*out*_ and *T*_*out*_. Each data point in the time courses represents the average of five consecutive strides and shaded areas around the data points represent the standard errors. Bar plots indicate the mean average behavior in the epochs of interest (indicated with the black rectangles), the gray dots indicate values for individual subjects, and vertical black lines are standard errors. Horizontal lines between bars illustrate significant differences between groups (*p* < 0.025). There was a significant group effect on *S*_*out*_ and *T*_*out*_. **A)** Steady States values of *T*_*out*_ and *S*_*out*_: We found significant group differences in *S*_*out*_’s and *T*_*out*_’s steady state. Colored asterisks indicate that the mean steady state for that group is significantly different from zero (*p* < 0.0125). **B)** After-effect values of *T*_*out*_ and *S*_*out*_: We found a significant group difference in *S*_*out*_’s after-effects. Colored asterisks indicate that the mean after-effect for that group is significantly different from zero (*p* < 0.0125).

The possible interdependence in space and time was further supported by the analysis of spatial and temporal features known to be adapted by the split-belt task, but not directly targeted by our feedback. Namely, the temporal feedback also modified the Adaptation and Post-adaptation time courses of the legs’ orientation asymmetry, quantified by *S*_*A*_, which is a spatial measure related to step position. Note that the time courses of *S*_*A*_ for the temporal feedback group (yellow trace) and control group (red trace) do not overlap during Adaptation and Post-adaptation (left panel Figure 5A and 5B). In contrast, the time courses of double support asymmetry (*T*_*A*_) were not altered by the temporal feedback, as shown by the overlap of *T*_*A*_ values during Adaptation and Post-adaptation of the temporal feedback and control groups (right panel Figure 5A and 5B). Consistently, we found a group effect in *S*_*A*_ (*p* = 0.0029) and a non-significant group (*p* = 0.8151) or group by epoch interaction (*p* = 0.3189)) in *T*_*A*_. *Post-hoc* analyses revealed that these effects were driven by group differences in *S*_*A*_’s steady state (*T → S*_*A*_ : *p* = 0.0138) and *S*_*A*_’s after-effects (*T → S*_*A*_ : *p* = 0.0163). Suprisingly, we did not find differences on *T*_*A*_’s steady state and after-effects, which we expected given the relation between *T*_*A*_ and the temporal measure (*T*_*out*_) directly altered with the temporal feedback. Thus, after-effects in *S*_*A*_ and *T*_*A*_ were significantly different from zero in all groups (control group: *S*_*A*_ : *p* = 0.0009 and *T*_*A*_ : *p* = 0.0044; temporal feedback group: *S*_*A*_ : *p* = 0.0080 and *T*_*A*_ : *p* = 0.0009), but only those of *S*_*A*_ were reduced in the temporal feedback group compared to controls. In sum, these results indicate that temporal feedback did not have a ubiquitous effect in all gait parameters, but it did alter the adaptation and recalibration of the legs’ orientation, which also characterizes the spatial control of the limb in locomotion.

**Figure 5:**
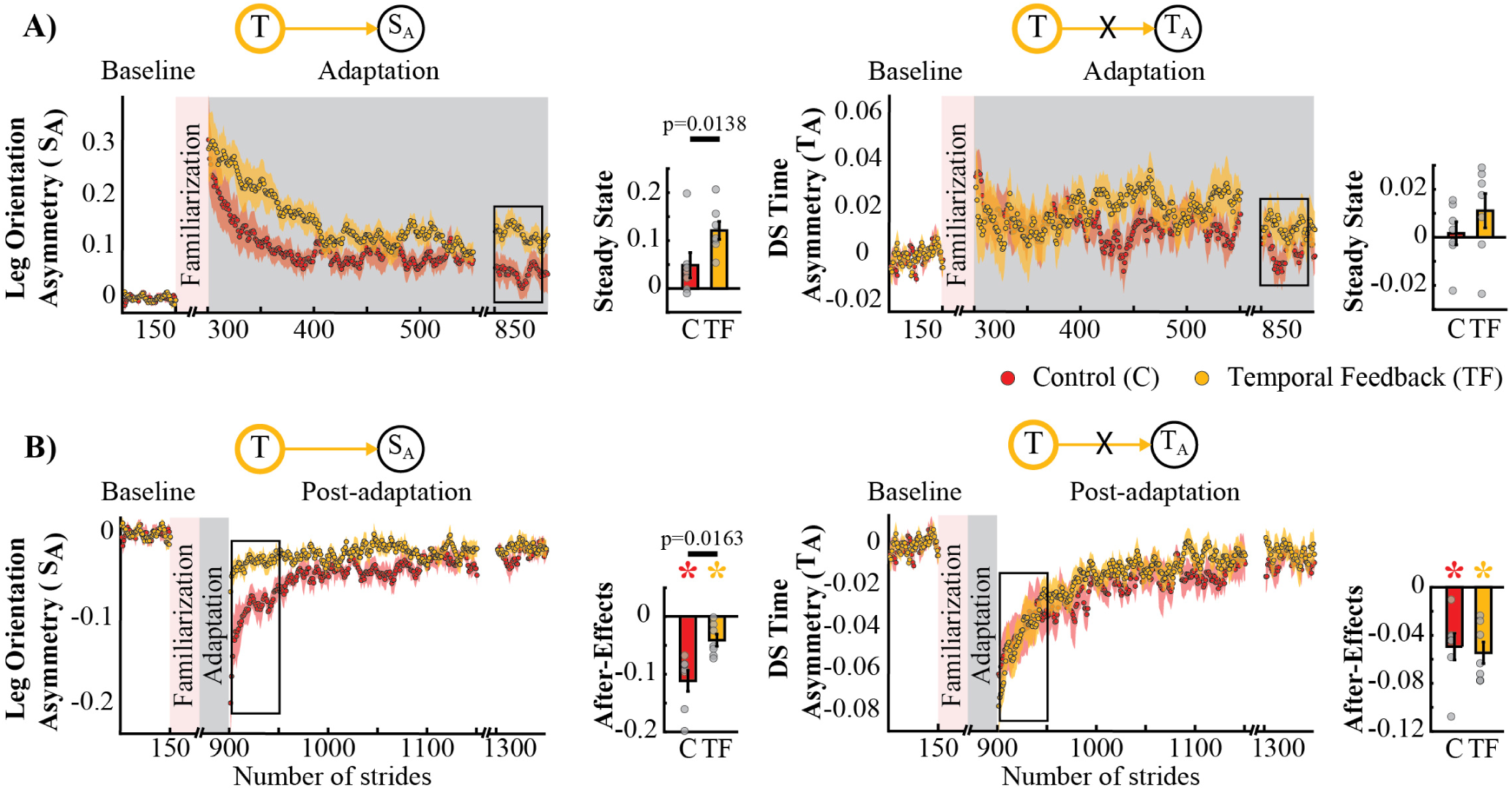
Adaptation and Post-adaptation for the adaptive but non-targeted parameters *S*_*A*_ (leg orientation asymmetry) and *T*_*A*_ (double support time asymmetry) in the temporal feedback and control groups. Stride-by-stride time courses show the effect of altering step times in the Adaptation (Panel A) and Post-adaptation (Panel B) of *S*_*A*_ and *T*_*A*_. Each data point in the time courses represents the average of five consecutive strides and shaded areas around the data points represent the standard errors. Bar plots indicate the mean average behavior in the epochs of interest (indicated with the black rectangles), the gray dots indicate values for individual subjects, and vertical black lines are standard errors. Horizontal lines between bars illustrate significant differences between groups (*p* < 0.025). There was a significant group effect in *S*_*A*_, but no in *T*_*A*_. **A)** Steady State values of *T*_*A*_ and *S*_*A*_: The significant group effect on *S*_*A*_ was driven by differences between the temporal feedback and control group in the non-targeted spatial motor output (adaptive motor output). **B)** After-Effects of *T*_*A*_ and *S*_*A*_: We found a significant group difference in *S*_*A*_. Colored asterisks indicate after-effect values are significantly different from zero (*p <* 0.025) according to *post-hoc* analysis.

### Temporal feedback modified the split-belt task to a greater extent than the spatial feedback

Surprisingly, temporal feedback altered the difference in stance times between the legs (*T*_*nA*_), whereas the spatial feedback did not. This was unexpected given previous literature indicating that *S*_*nA*_ and *T*_*nA*_ do not change as subjects walk in the split-belt environment (Malone et al., 2012; Reisman et al., 2005; Yokoyama et al., 2018). Thus, we anticipated that either type of feedback (spatial or temporal) would not alter these “non-adaptive” gait features. Qualitatively, we observed that this was the case for the spatial (*S*_*nA*_), but not for the temporal (*T*_*nA*_) “non-adaptive” parameter (Figure 6A). Note that *S*_*nA*_ has the same time course for both groups, whereas *T*_*nA*_ has a different time course for the control group (red trace) and the temporal feedback group (yellow trace). Consistently, we found a significant group effect (*p* = 0.0030) and group by epoch interaction (*p* = 0.0047) in *T*_*nA*_, whereas a non-significant group (*p* = 0.3860) or group by epoch interaction effect (*p* = 0.3719) in *S*_*nA*_. *Post-hoc* analysis revealed that the temporal feedback group reached a significantly lower steady state when compared to the control group (*T → T*_*nA*_ : *p <* 0.0001). Conversely, the spatial feedback group exhibited the non-adaptive behavior of these parameters *S*_*nA*_ and *T*_*nA*_ that we anticipated. Namely, the time courses of *S*_*nA*_ (Figure 6B, left panel) and *T*_*nA*_ (Figure 6B, right panel) were overlapping in these two groups. This similarity is subtantiated by the the non-significant group effect (*S*_*nA*_ : *p* = 0.2338 and *T*_*nA*_ : *p* = 0.3002) or group by epoch interaction (*S*_*nA*_ : *p* = 0.7452 and *T*_*nA*_ : *p* = 0.8163) in the non-adaptive spatial and temporal parameter. In sum, feedback modifying the adaptation of spatial and temporal gait features had a distinct effect on “non-adaptive” temporal parameters thought to only depend on the speed difference between the legs in the split-belt task.

**Figure 6:**
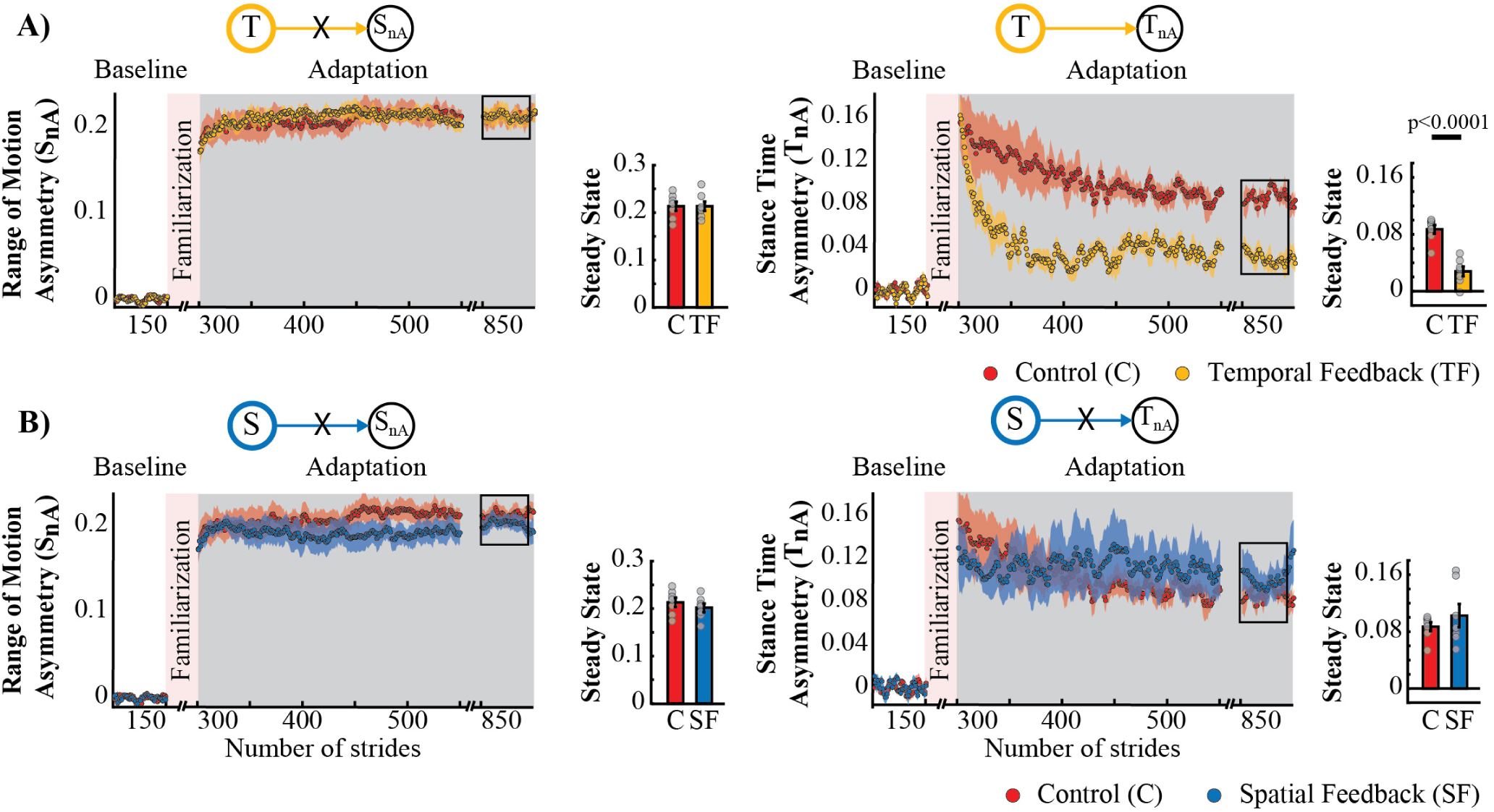
Adaptation of *S*_*nA*_ and *T*_*nA*_ measures that are non-adaptive and non-targeted parameters in temporal feedback and control group (Panel A) and spatial feedback and control group (Panel B). Stride-by-stride time courses show the effect of altering the step times or step positions on “non-adaptive” temporal and spatial measures (*S*_*nA*_ and *T*_*nA*_) during Adaptation. Each data point in the time courses represents the average of five consecutive strides and shaded areas around the data points represent the standard errors. Bar plots indicate the mean average behavior in the epochs of interest (indicated with the black rectangles), the gray dots indicate values for individual subjects, and vertical black lines are standard errors. Horizontal lines between bars illustrate significant differences between groups (*p <* 0.025). **A)** Steady State values of *T*_*nA*_ and *S*_*nA*_: We found a significant group effect and group by epoch interaction driven by differences between the temporal feedback and control group in the non-targeted temporal motor output (adaptive motor output). **B)** Steady State values of *S*_*nA*_ and *T*_*nA*_: We did not find a significant group effect or group by epoch interaction for the spatial feedback and control group in the parameters of interest.

## 4 DISCUSSION

### 4.1 Summary

Our study confirms previous results suggesting that there are internal representations of space and time for predictive control of movement. We replicated previous results showing that altering the recalibration in the spatial domain does not impact the temporal domain. However, we also observed that the opposite was not true. That is, explicitly reducing the recalibration in the temporal domain altered movement control in space, suggesting some level of interdependence between these two domains. Interestingly, double support asymmetry was consistently corrected across the distinct spatio-temporal perturbations that subjects experienced, whereas spatial asymmetries were not. This indicates that correcting asymmetries in space and time is prioritized differently by the motor system. Our results are of translational interest because clinical populations often have greater deficits in either the spatial or the temporal control of the limb and our findings suggest that they may not be treated in isolation.

### 4.2 Separate representations for predictive control of movements in space and time

We find that adaptation of movements to a novel walking situation results in the recalibration of internal representations for predictive control of locomotion; which are expressed as robust after-effects in temporal and spatial movement features. This is consistent with the idea that the motor system forms internal representations of space (Marigold and Drew, 2017) and time (Avraham et al., 2017; Breska and Ivry, 2018; Drew and Marigold, 2015) for predictive motor control. Several behavioral studies suggest separate recalibration of these internal representations of space and time in locomotion because spatial and timing measures exhibit different adaptation rates in the mature motor system (Malone and Bastian, 2010; Darmohray et al., 2019) throughout development (Vasudevan et al., 2011; Patrick et al., 2014) or healthy aging (Sombric et al., 2017). Spatial and temporal recalibration also have distinct generalization patterns across walking environments (Torres-Oviedo and Bastian, 2010; Mariscal et al., 2018) and most importantly, altering the adaptation of spatial features does not modify the adaptation and recalibration of temporal ones, as shown by us and others (Malone et al., 2012; Long et al., 2016). This idea of separate representations of space and time in locomotion is also supported by clinical and neurophysiological studies indicating that different neural structures might contribute to the control (Rybak et al., 2006; Lafreniere-Roula and McCrea, 2005) and adaptation (Vasudevan et al., 2011; Choi et al., 2009; Statton et al., 2018) of the spatial and temporal control of the limb in locomotion.

### 4.3 Hierarchic control of timing leads to interdependent adaptation of movements in space and time

Nonetheless, we also found that explicit control of step timing modifies the adaptation and recalibration of movements in space. This result directly contradicts the dissociable adaptation of spatial and temporal features upon explicitly modifying the adaptation of step position (spatial parameter) (Malone et al., 2012; Long et al., 2016). We find two possible explanations to reconcile these findings. First, there might be a hierarchical relationship between the spatial and temporal control of the limb, such that timing cannot be manipulated without obstructing the adaptation of spatial features. We believe that this type of hierarchical organization is not exclusive to explicit control, but it is also applicable to implicit control of the limb in space and time. This is supported by a recent study indicating that lesions to interpose cerebellar nuclei altering the adaptation of double support asymmetry (temporal parameter) also reduced the after-effects of spatial features (Darmohray et al., 2019), whereas the recalibration of spatial features can be halted without modifying the temporal ones (Darmohray et al., 2019). Future studies are needed to determine if similar results would be observed in human bipedal locomotion. This type of hierarchical organization suggests that the execution of spatial and temporal control of the limb can be encoded by separate interneuronal networks (Rybak et al., 2006; Lafreniere-Roula and McCrea, 2005), but the volitional recruitment of those networks cannot occur in isolation. Second, it is possible that the observed interdependence arose as a byproduct of how we tested it. Namely, it remains an open question if our findings result from altering step time, or similar interdependence would be observed if we had manipulated other temporal measures, such as double support asymmetry. More specifically, our feedback on step time inadvertently reduced the stance time asymmetry associated to split-belt walking. The stance time asymmetry is thought to be critical for forcing subjects to adjust their gait during split-belt walking (Reisman et al., 2005). Therefore, subjects in the temporal feedback group might have reduced the adaptation of spatial parameters because the “perturbation” inducing their update was reduced. In sum, future work is needed to determine the generality of temporal measures influencing spatial ones, however our study provides initial evidence for interdependence.

### 4.4 Relevance of double support symmetry over spatial asymmetries

We demonstrated that double support symmetry (i.e., *T*_*A*_) is recovered in all groups, regardless of the task. This is in accordance with multiple observations that individuals consistently reduce double support asymmetries induced by split-belt walking since very early age (Patrick et al., 2014) or after lesions to cerebral (Reisman et al., 2007) or cerebellar regions (Vasudevan et al., 2011). Only children with hemispherectomies, where half of the cerebrum is missing, do not correct double support asymmetry when this is augmented (Choi et al., 2009). The adaptation and after-effects of double support were surprising to us because previous work showed that halting the adaptation of step position (*S*_*out*_ ≈ 0) limited the correction of spatial errors (defined as *S*_*A*_) (Malone et al., 2012). In an analogous manner, we anticipated that preventing the adaptation of step times (*T*_*out*_ ≈ 0) during split-belt walking was going to limit the adaptation of double support asymmetry (i.e., temporal error (Malone et al., 2012)). However, we observed that individuals prioritize differently the correction of spatial and temporal asymmetries: they minimize temporal asymmetries, but not spatial ones. This might be because double support time is the transition period when the body mass is transferred from one leg to the other, which is demanding in terms of energy expenditure (Perry, 1992). Therefore, double support symmetry might be critical for efficient body transfer between the limbs (Kuo et al., 2005; Ruina et al., 2005). Taken together our results suggests that the motor system prioritizes the maintenance of double support symmetry, which might be critical for balance control in bipedal locomotion.

### 4.5 Explicit vs. implicit processes in locomotor adaptation

Our study contributes to recent efforts to unveil the potential interaction between explicit corrections and implicit sensorimotor recalibration in locomotion (Statton et al., 2016; Roemmich et al., 2016; Long et al., 2016; Malone et al., 2012; Maeda et al., 2017). Interestingly, we found that preventing foot adjustments during split-belt walking significantly reduced post-adaptation effects compared to the control group. This was also observed when using explicit corrections to reduce the adjustment of foot placement in response to a 2:1 speed belt ratio (Malone et al., 2012) but not in response to a larger 3:1 speed belt ratio (Long et al., 2016). Notably, after-effects following the 3:1 perturbation were equally large with or without explicit corrections during the split condition (Long et al., 2016). One interpretation for these results is that the implicit sensorimotor adaptation in walking is scaled with perturbation magnitude. Thus, explicit corrections preventing foot adjustments in the split condition will have a lesser impact on after-effects induced by large perturbations. This interpretation is consistent with the proportional relation between perturbation size and after-effects upon experiencing unexpected constant forces (Yokoyama et al., 2018; Green et al., 2010; Torres-Oviedo and Bastian, 2012), contrasting the fixed amount of implicit sensorimotor recalibration upon visuomotor perturbations (Kim et al., 2018).

### 4.6 Study implications

We provide a novel approach for manipulating stance time, which is a major deficit in stroke survivors (Patterson et al., 2008). It would be interesting to determine if this type of feedback overground or on a regular treadmill could lead to gait improvements post-stroke as those induced by split-belt walking (Reisman et al., 2013; Lewek et al., 2018). Our results also indicate that manipulating the adaptation of movements in the temporal domain alters movements in the spatial domain, suggesting that spatial and temporal deficits in individuals with cortical lesions (Finley et al., 2015; Malone and Bastian, 2014) cannot be treated in complete isolation. Only the correction of timing asymmetries through error-based sensorimotor adaptation could occur while preventing the adaptation of spatial ones, as we did in the spatial feedback group. However, the opposite is not possible, at least with the temporal feedback task that we used.

## AUTHOR CONTRIBUTIONS

M.G. and N.V. equally contributed to data acquisition and processing (Gonzalez-Rubio et al., 2019). They also contributed in the interpretation of the data and final approval of the version to be published, and agreement to be accountable for all aspects of the work. G.T-O. contributions include conception and design of the work, analysis of the data, writing a complete draft of the manuscript, revising work for important intellectual content, final approval of the version to be published, and agreement to be accountable for all aspects of the work.

## FUNDING

The project was funded by National Science Foundation (NSF1535036), and American Heart Assosiation (AHA 15SDG25710041).

### ACKNOWLEDGMENTS

The authors acknowledge the valuable input from Pablo Iturralde and Carly Sombric.

## DATA AVAILABILITY STATEMENT

The datasets generated and analyzed for this study can be found in the *Figshare* repository [https://figshare.com/articles/ExplicitTemporal_SpatialModulations_mat/8145962]

